# ChiMera: An easy to use pipeline for Bacterial Genome Based Metabolic Network Reconstruction, Evaluation and Visualization

**DOI:** 10.1101/2021.11.30.470608

**Authors:** Gustavo Tamasco, Ricardo R. da Silva, Rafael Silva-Rocha

## Abstract

Several genome scale metabolic reconstruction tools have been developed in the last decades. They have helped to construct many metabolic models, which have contributed to a variety of fields, e.g., genetic engineering, drug discovery, prediction of phenotypes, and other model-driven discoveries. However, the use of these programs requires a higher level of bioinformatic skills, and most of them are not scalable for multiple genomes. Moreover, the functionalities required to build models are generally scattered through multiple tools, requiring knowledge of their utilization.

Here, we present ChiMera, which combines the most efficient tools in model reconstruction, prediction, and visualization. ChiMera uses CarveMe top-down approach based on genomic evidence to prune a global model with a high level of curation, generating a draft genome able to produce growth predictions using flux balance analysis for gram-positive and gram-negative bacteria. ChiMera also contains two modules of visualization implemented, predefined and universal. The first generates maps for the most important pathways, e.g., core-metabolism, fatty acid oxidation and biosynthesis, nucleotides and amino acids biosynthesis, glycolysis, and others. The second module produces a genome-wide metabolic map, which can be used to harvest KEGG pathway information for each compound in the model. A module of gene essentiality and knockout is also present. Overall, ChiMera combines model creation, gap-filling, FBA and metabolic visualization to create a simulation ready genome-scale model, helping genetic engineering projects, prediction of phenotypes, and other model-driven discoveries in a friendly manner.

## Introduction

Genome-Scale Metabolic Reconstruction (GSMR) is an essential tool in System Biology (1). The use of GSMR helped researchers over the last 30 years to understand microbial evolution, network interaction, genetic engineering, drug discovery, prediction of phenotypes, and model-driven discoveries (2). However, the production of a precise model is very complex and time-consuming, requiring several steps. The process starts with genome annotation and the prediction of possible metabolite and reaction candidates, which creates an initial database to build the draft model. Several rounds of manual curation and evaluation of the present genes, reactions, and compounds are necessary to create a precise model. After this step, one needs to set the Objective Function of the model, e.g., biomass function, followed by the conversion to a mathematical formulation entitled Stoichiometric matrix (S-matrix), which is the core of Flux Balance Analysis (FBA) and growth predictions (3). Other steps, e.g., gap-filling and stoichiometric balance may also be necessary, increasing the complexity of the process.

Recently, several tools were developed to address some of these steps (4). CarveMe (5) is a command-line tool that deals with the initial phase of model creation and gap-filling. Cobrapy (6) can convert draft models into an S-matrix and perform FBA analysis using optimized algorithms. Escher (7) offers a fully customizable suite for pathway visualization. However, these tools require familiarity with command line interfaces and programming. They also have their peculiarities, demanding time and knowledge from users to perform the analysis. Therefore, the use of those tools by non-bioinformatics can be challenging, and the number of steps required to build initial models precludes their usage in large-scale projects (i.e., hundreds of genomes).

Here, we demonstrate a novel tool named ChiMera, which uses automation algorithms to compile the most used tools in literature in a user-friendly pipeline that does not require programming skills. ChiMera uses a genome as input (*.faa file), performing the model creation based on a highly curated universal model. That is used for FBA and growth predictions, knockout simulations, and pathway visualization (Figure 1). To evaluate ChiMera, we compared several aspects of model completion with manually curated models from the literature. We also compared the predicted growth values with experimental data to ensure the production of realistic values.

**Figure 1:**
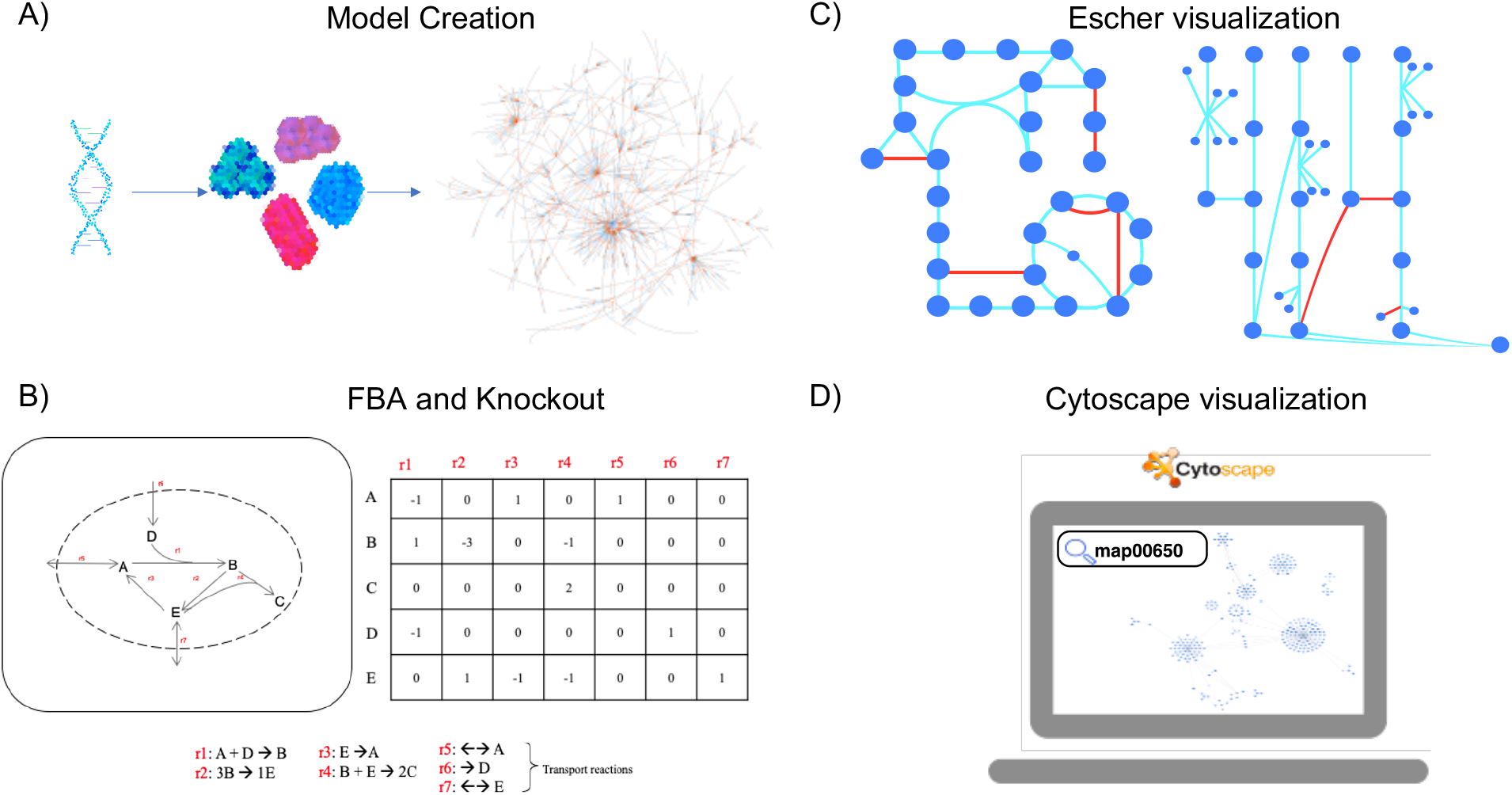
Processes automated by Chimera. Chimera uses automation to compile several tasks and create an organism specific model based on the genomic data. A) Model creation based on CarveMe pruning algorithm, users just need to provide a protein annotation file based on the genome, B) Model conversion to a Stoichiometric Matrix to perform Flux Balance Analysis and Gene and Reaction Knockouts. C) Representation of pre-defined Escher maps, reactions depicted in blue were detected in the model, red ones were absent. D) Pathway specific compounds detected using the Chimera translator module (add pathway information to compounds), here butanoate metabolism is depicted.

## Material and Methods

### General ChiMera structure

ChiMera uses automation algorithms to combine three main steps in GSMR, model creation and gap-filling, flux balance analysis, and pathway visualization. Along with a knockout prediction module based on FBA, which enables gene essentiality evaluation. ChiMera is modular as discussed in the results and compatible with further module expansions.

### Model Creation and gap-filling

We automated the utilization of CarveMe (v1.5.1) in the reconstruction module of ChiMera. The initial draft model is created based on the protein file (*.faa file) provided by the user. During the reconstruction process, ChiMera also performs a gap-filling based on the medium definition, using genomic evidence to ensure that the model will produce growth under the given conditions. CarveMe uses a top-to-bottom approach in a reference pre-built manually curated universal model. It applies a pruning algorithm that removes reactions not supported by the genomic evidence, generating an organism-specific model based on highly curated data (5).

### S-matrix construction and initial FBA

We used COBRApy (v0.22.1) to convert the initial draft into an S-matrix and perform an FBA analysis. The tool is also the core behind the ChiMera automated pipeline to perform gene and reaction knockouts (6).

### Visualization of the metabolic maps

We developed *in-house* algorithms to automate the generation of metabolic maps based on Escher (v1.7.3). Ten predefined pathway maps are pre-loaded in this module. Users can also provide custom JSON maps of desired pathways to check if they are present in the target organism. The pipeline uses the model data to evaluate reactions and compounds present in the organism, creating customizable HTML maps that can be edited by the user (7).

We also developed a module that couples COBRApy data with PSAMM (v1.1.2), converting the model to a graphical representation (8). Users can also convert the BiGG ids to KEGG ids and collect pathway information for compounds. This module produces a file that can be loaded into Cytoscape, being compatible with all the well-established plugins of the tool (9).

### Genome Selection

We selected two well-studied organisms representing Gram-negative bacteria, *Pseudomonas putida* KT2440 and *Escherichia coli. Bacillus subtilis* was selected to represent Gram-positive bacteria. Protein files were downloaded from NCBI under the accession number <u>NC_002947.4</u>, <u>NZ_CP020543.1</u> and <u>AL009126.3</u>. We compared the models generated by ChiMera to manually curated models from the BiGG database, iJN746, iEC1344_C and iYO844 respectively.

### Model evaluation

We performed basic tests to check the correctness of the models produced by ChiMera using MEMOTE which benchmarks the model using consensus tests based on model annotation, biomass composition and stoichiometry. The biomass production, presence of fundamental metabolites in the biomass function, stoichiometric balance and inconsistency were some of the parameters compared to highly curated models (10).

We performed a gene essentiality benchmark to assay the effect of a single-gene deletion. The media composition was defined as M9 minimal medium for all the organisms. To calculate the performance metrics, we measured the ChimMera’s ability to correctly assign a gene as non-essential or essential. Predicted outcomes were compared with experimental data (11–13).

### Performance metrics

We used 6 different performance metrics to compare the genes essentiality predictions from ChiMera to highly curated models.

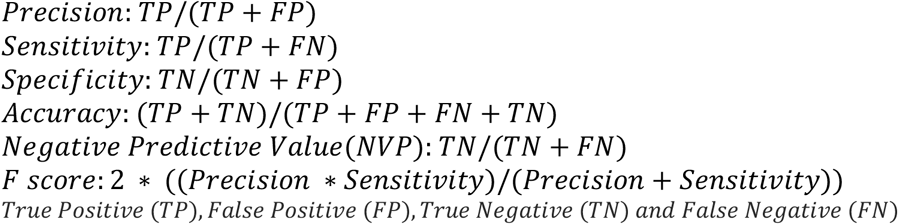

### ChiMera environment and user interface

ChiMera is a portable command-line-based tool. The source code, along with complete documentation of its utilization and examples of inputs are available (https://github.com/tamascogustavo/chimera) (14).

## Results

### Key Capabilities of ChiMera

ChiMera was implemented in python v3.7, and its dependencies are freely available. There are four main functionalities: Model Creation, Flux Balance Analysis and Growth prediction, Metabolism Visualization and Knockout evaluation (Figure 2). ChiMera relies on CarveMe to create an organism-specific model. A curated model is pruned to produce a draft model containing thermodynamic balanced reactions and elemental balanced compounds, which interact across three compartments, cytosol, periplasm, and extracellular. During the reconstruction, the user can select from up to six different mediums, and perform a gap-filling based on the genomic evidence to ensure that the organism can grow under the provided circumstances

**Figure 2:**
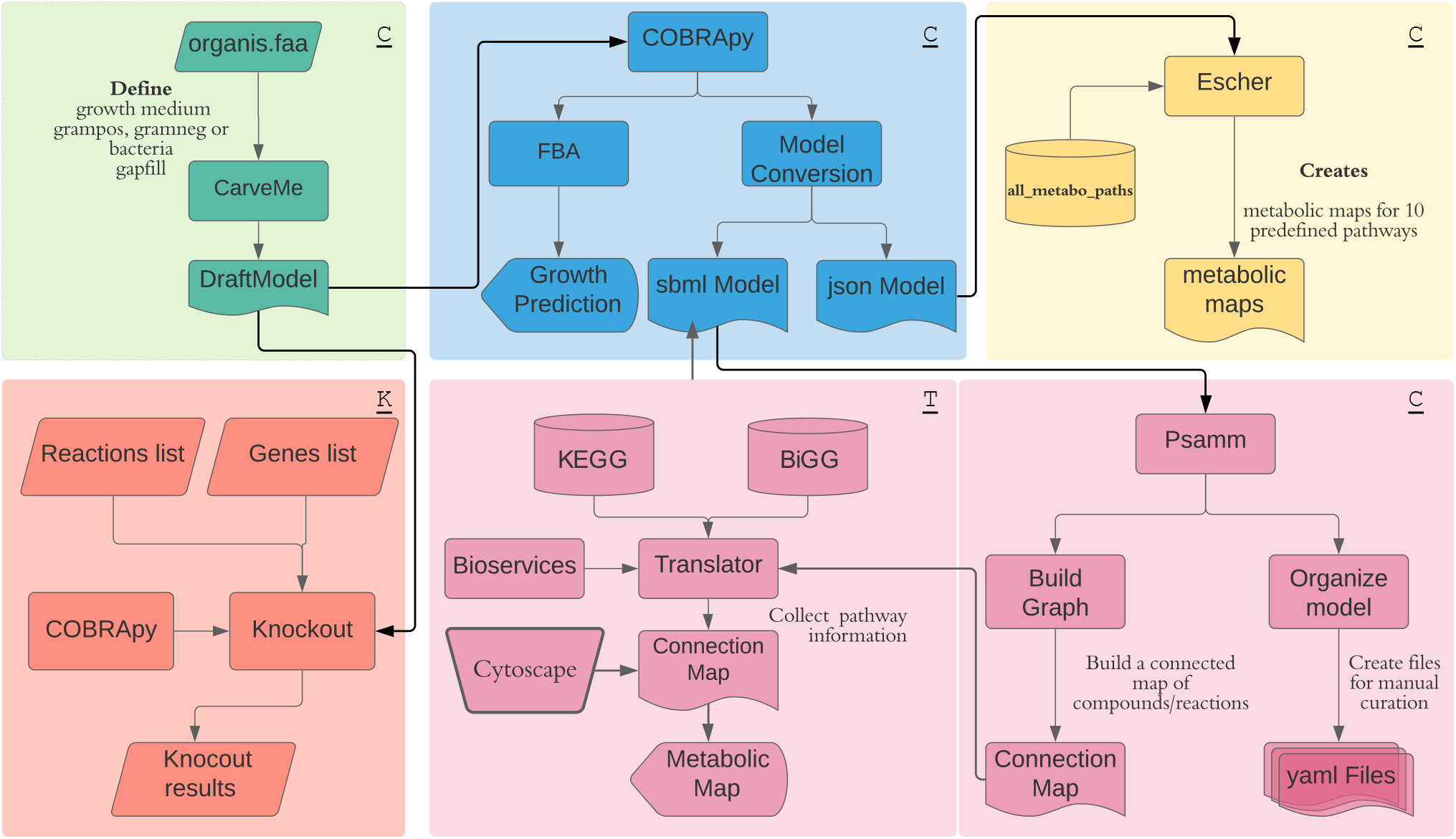
Flow chart of Chimera processes. Chimera has 3 submodules that can be used separately. The ones signed with “<u>C”</u> are part of the core module, which performs model creation, evaluation and creation of visualization files. The “<u>T</u>” represents the translator module that adds KEEG pathway information to the compounds. The file can be loaded into Cytoscape for visualization. The “<u>K</u>” represents the knockout module, which performs gene and reaction knockouts.

The organism-specific model is automatically converted to a S-matix, using COBRApy. The biomass reaction is set as the Objective Function for FBA. The flux values for each metabolite in the biomass function are informed to the user, along with the normalized growth value (1 mmol / g[CDW] / h) and basic model statistics (Supplementary Figure 1). Subsequently, the model is converted to a json format, used to produce predefined metabolic maps based on Escher. They include carbohydrate, central carbon, fatty acid oxidation and biosynthesis, glycolysis, inositol, and tryptophan metabolism (Figure 1C). The model is also converted to yaml format, which is used by PSAMM *findprimalpairs* algorithm to break down the GSMR into connections between compounds (nodes) and reactions (edges).

The output of PSAMM can be directly loaded into Cytoscape, producing a visualization of the entire reconstruction. Users can also use the ChiMera translator submodule, to add pathway information to the file, enabling a targeted search of pathways in Cytoscape (Figure 1 D).

To allow ChiMera’s flexibility and modularity, users can also provide a pre-built model with the protein file, which should hold the same prefix, directly performing FBA analysis and construction of the pathway maps. Documentation is provided to ensure that the annotations of the model or the presence of extra compartments are compatible with PSAMM, to generate the Cytoscape compatible file.

The knockout module is dependent on COBRApy functionality. Here, we implemented a function that enables the user to provide a file (.txt) containing a list of genes or reactions to be silenced. This module can perform single or double targeted deletions. The user can also perform gene essentiality analysis for the whole model, identifying the impact of silencing on the growth under given circumstances.

### Comparison with manually curated models

Models created by ChiMera had a higher presence of exclusive reactions and compounds except for *E. coli*. Comparing shared compounds between models, around 68% of metabolites are present in both scenarios. For reactions, the average is lower, reaching only 60% of shared reactions (Figure 3).

**Figure 3:**
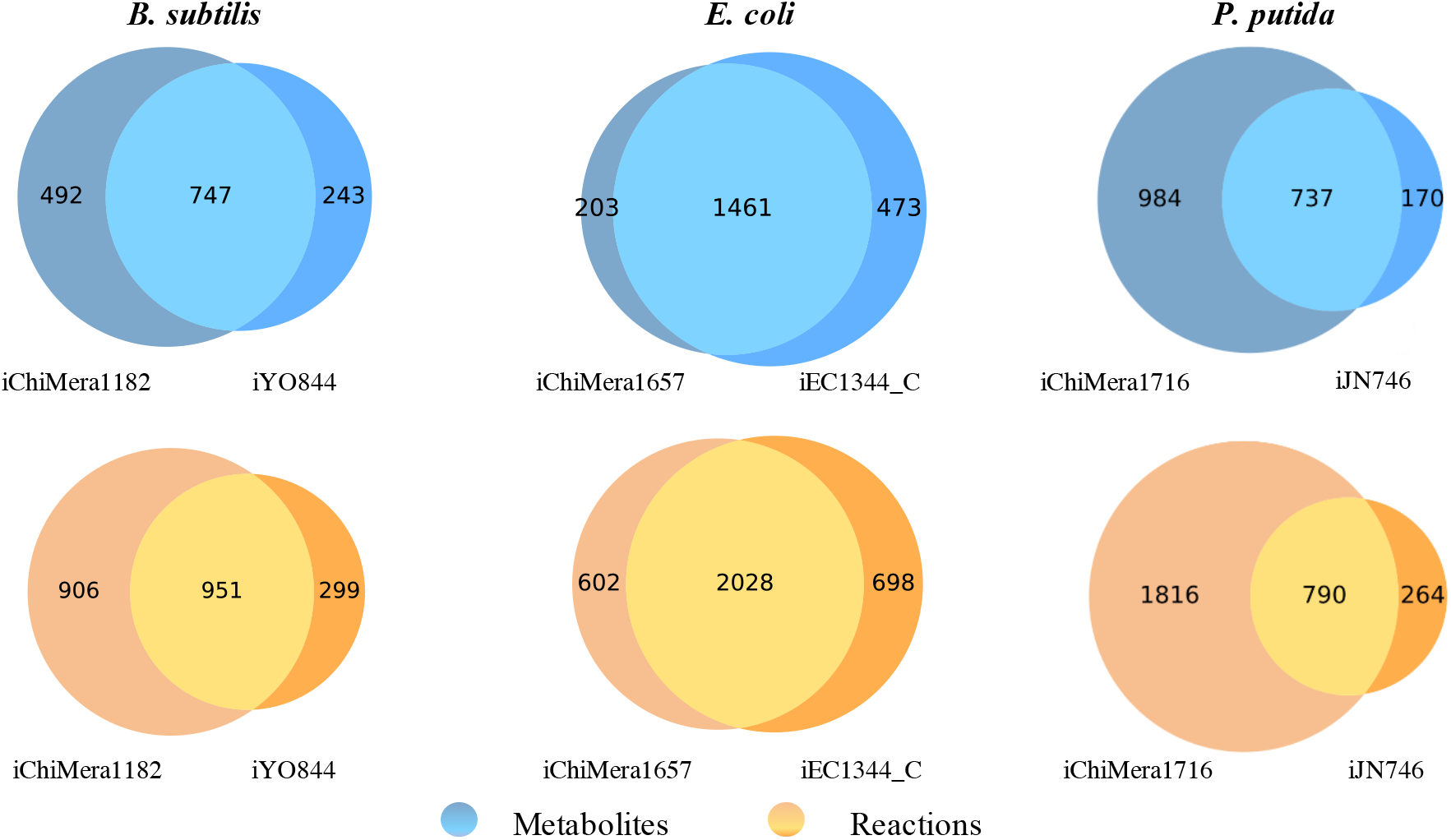
Venn Diagram of Reactions and Metabolites sets. Reactions and compound sets from Chimera and manually curated models were compared to identify the intersection of the model composition. Model exclusive information is also depicted.

The overall MEMOTE score of ChiMera models is lower than the manually curated ones, due to the lack of gene annotation in the final draft. However, ChiMera has a lower number of blocked reactions, orphan and dead-end metabolites, which play a role in the accuracy of model predictions. Moreover, curated models had a higher presence of missing essential precursors in the Biomass Function, which can lead to unrealistic growth predictions (Table 1).

**Table 1:**
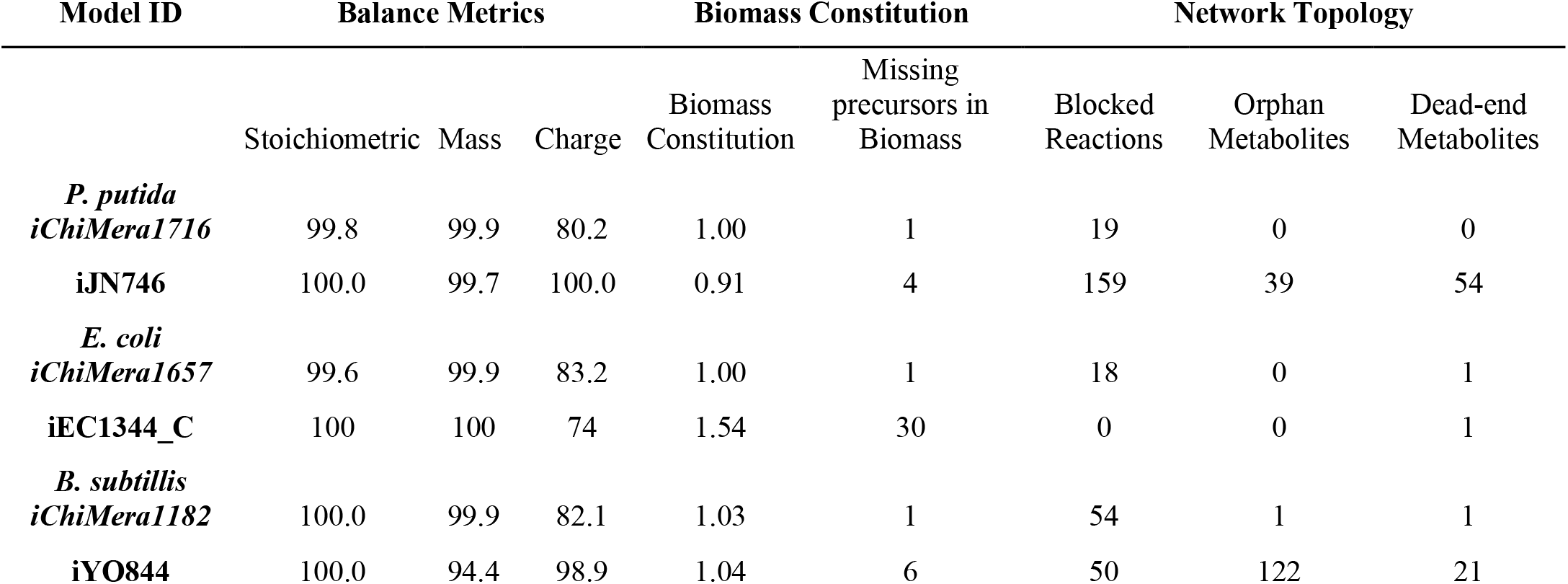
MEMOTE evaluation metrics. Parameters that can influence the precision of the predictions were selected to assay Chimera and BiGG curated models. Values in the range of 1±10-3 in Biomass Constitution are necessary to indicate a realistic biomass function.

ChiMera produces models in different formats, SBML3, JSON and YAML. These models can be used as input for Merlin (15) or PSAMM (8), enabling advanced users to perform manual curation of their models. The gene essentiality predictive metrics were higher in manually curated models. For *E. coli*, the iEC1344_C had a perfect prediction for the tested dataset. The ChiMera model was slightly outperformed in all predictive metrics (Figure 4A). For *B. subtilis*, the iYO844 performed better than ChiMera model, which had higher error rates in Sensitivity and Negative Predictive Value, due to the higher presence of False Negative predictions (Figure 4B). A similar trend was observed for *E. coli*. The *P. putida* model outperformed iJN746 by a small margin. The improvements were in Precision and Specificity, indicating less detection of False Positive values in *P. putida* automated model (Figure 4C).

**Figure 4:**
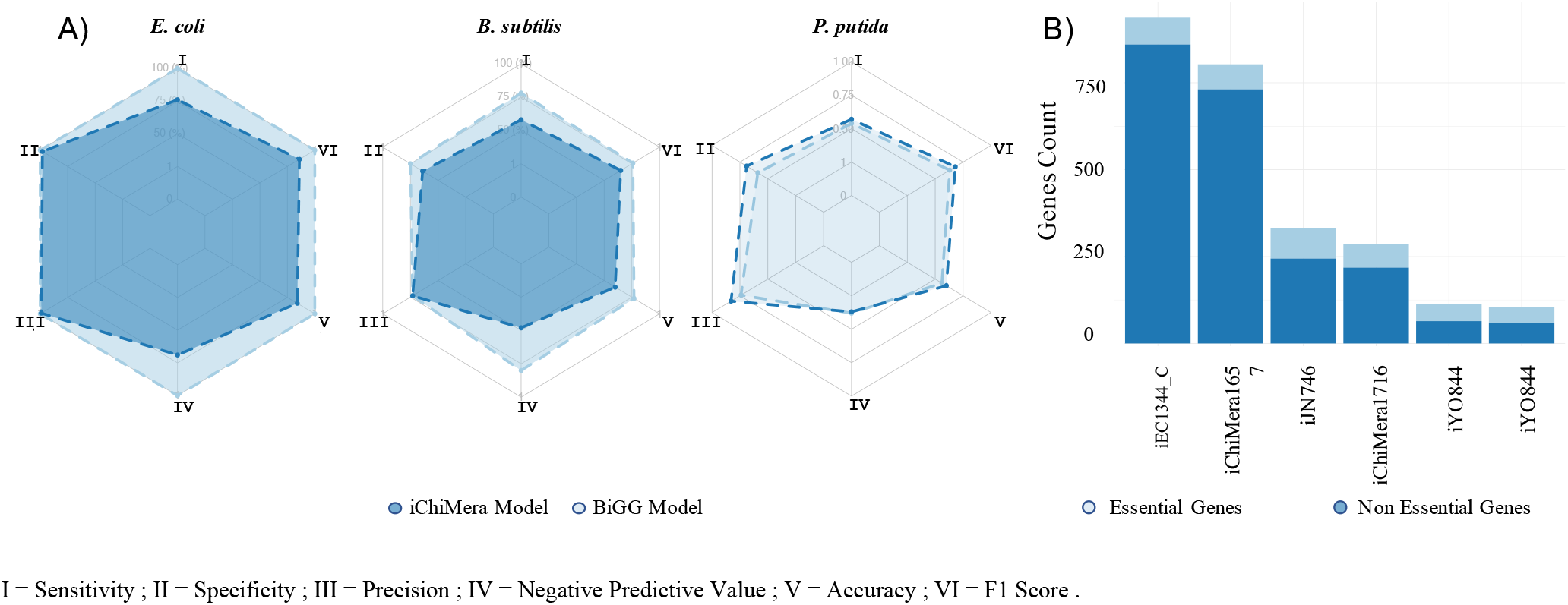
Gene essentiality metrics. Six metrics were selected to compare prediction capability from Chimera and manually curated models. A) Radarplot of Gene Essentiality metrics. Non essential genes were used as true positive and essential genes as true negative. B) Stacked bar plot of gene essentiality classification according to presence in the genome.

## Discussion

We introduce ChiMera, an automated, well documented and easy to use command line tool that enables non-bioinformaticians to produce Genome-Scale Metabolic Reconstructions. The models can be used to explore the metabolic potential of the target organism. Gene essentiality modules can help researchers to understand the behavior of the organism under disturbed experimental conditions. The visualization modules facilitate the exploration of essential pathways, as well as the identification of unique pathways for non-model organisms. In sync, the information provided by ChiMera assists researchers in understanding non-model organism metabolism or even developing engineering approaches for model ones.

ChiMera has similar ambitions to AuReMe and Merlin. These tools offer a custom workspace for the user, hence facilitating the construction of Genome-Scale Models. AuReMe has its own data structure based on PADMet, and focuses on traceability of the reconstruction process, performing at its best if highly curated models are available (16). There are several steps that the user can process, but it lacks visualization and knockout modules (Supp Table 1). Its performance was comparable to CarveMe in model creation (4).

Merlin offers a vast workspace for its users. Its graphical interface allows users to re-annotate genomes using BLAST or HMMER, and also integrate data from NCBI and KEGG to its draft model (15). This tool is preferable for those focusing on manual curation of single organisms with expertise in metabolic engineering and model creation (4).

ChiMera inherits some pros and cons from CarveMe. Generating networks that share coverage of reactions and metabolites above 60% compared to highly curated models (Figure 3). Which shows a great potential for first model draft, prior to manual curation. The ready to simulate models of ChiMera are also valuable assets for those working with hundreds of genomes due to the easiness and speed of a draft construction, enabling researchers to evaluate multiple candidate models and choosing the best option for a manual curation if needed.

Here we demonstrate that manually curated models mostly had higher prediction capabilities when compared to models created by ChiMera. However, in some cases, ChiMera models outperform curated models, and when outcompeted the scores were closely related to them (Figure 4a). The visualization components allow the user to easily browse and improve the model, therefore, ChiMera can also be used in semi-automated capacity, shortening the cycles of manual curation. Thereby, ChiMera can be a valuable asset for investigating non-model organisms, providing knowledge of its metabolism, asslowing scale reconstruction and simulation of metabolic models starting from annotated genomes. ChiMera can even be used to explore gene essentiality aspects in the genomes of interest. All included in a friendly pipeline, fully compatible with further expansion of its modules, which show the general utility of the tool to the scientific community.

The authors declare that the research was conducted in the absence of any commercial or financial relationships that could be construed as a potential conflict of interest.

## Author Contributions

R.D.S and RS-R contributed to the review of the paper and ideas of design. GT was responsible for the study design and implementation, script creation, data analysis and wrote the draft of this manuscript. All authors contributed to manuscript revision, read, and approved the submitted version

## Acknowledgments

The authors would like to thank Mateus Gonçalves, Daniela Vicentini and Beatriz Bergamo for testing ChiMera and for all the feedbacks. We also would like to thanks Maria Eugenia for a final review of this manuscript.

## Supplementary Material

The Supplementary Material for this article can be found at the end of this document.

## Data Availability Statement

ChiMera is freely available at: https://github.com/tamascogustavo/chimera

