## Supplemental Table 1 Figure 1 for "ChiMera: An easy to use pipeline for Bacterial Genome Based Metabolic Network Reconstruction, Evaluation and Visualization"

Supplementary Material

**Supplementary Figure 1:** Chimera core module growth prediction output. The core module shows in the user screen the growth rate given the conditions provided to Chimera.

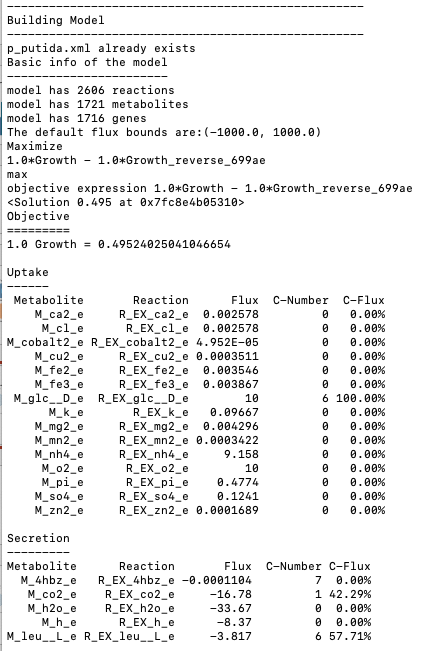
**Supplementary Table 1:** List of Genome Scale Metabolic Reconstruction Tools and their functionality.

| **Reconstruction Tool** | **Mapping Method** | **Reaction origin** | **Associated Database** | **Version** | **Visualization** | **Knockout** | **Type of Software** |
| --- | --- | --- | --- | --- | --- | --- | --- |
| AureMe | Pantograph (Inparanoid and OrthoMCL) | Template model(s) | BiGG-MetaCyc | 1.2.4 | No | No | Command line |
| CarveMe | Diamond, eggNOG-mappera | Template model | BiGG | 1.5.1 | No | No | Command line |
| ChiMeRa | Diamond | Template model  User defined model | BiGG | 1.0 | Yes | Yes | Command line |
| Merlin | Mapping from annotation with BLAST or HMMER | Template model(s) | KEGG | 4.0 | Yes | Yes | Stand Alone Interface |
| ModelSEED | Annotation ontology map from RAST data | Template model(s) | ModelSEED | 2.2-2.4 | Yes | No | Online service |
| Pathway Tools | Pathologic | Database | MetaCyc | 22.0 | Yes | Yes* | Stand Alone Interface |
| RAVEN | Autograph-type method from BLASTP and Bidirectional BLASTPb | Database | KEGG-MEtaCyc | 2.0.1 | Yes | No | Command line |

*means that the tool has dependency on external software to produce the knockouts, in this case Web-MetaFlux Modeling Tool.
